# Spatially organizing biochemistry: choosing a strategy to translate synthetic biology to the factory

**DOI:** 10.1101/202259

**Authors:** Christopher M. Jakobson, Danielle Tullman-Ercek, Niall M. Mangan

## Abstract

Natural biochemical systems are ubiquitously organized both in space and time. Engineering the spatial organization of biochemistry has emerged as a key theme of synthetic biology, with numerous technologies promising improved biosynthetic pathway performance. One strategy, however, may produce disparate results for different biosynthetic pathways. We propose a spatially resolved kinetic model to explore this fundamental design choice in systems and synthetic biology. We predict that two example biosynthetic pathways have distinct optimal organization strategies that vary based on pathway-dependent and cell-extrinsic factors. Moreover, we outline this design space in general as a function of kinetic and biophysical properties, as well as culture conditions. Our results suggest that organizing biosynthesis has the potential to substantially improve performance, but that choosing the appropriate strategy is key. The flexible mathematical framework we propose can be adapted to diverse biosynthetic pathways, and lays a foundation to rationally choose organization strategies for biosynthesis.

## Introduction

Synthetic biology traces its origins to the discovery of type II restriction endonucleases (Kelly and Smith, 1970; Smith and Welcox, 1970). These enzymes allowed the controlled assembly of novel genes, plasmids, and other nucleic acids, and precipitated the rapid spread of molecular cloning technology. Since then, synthetic biology has sought to exploit, adapt, and extend biological systems to benefit society by creating pharmaceuticals, fuels, gene therapies, drug delivery platforms, probiotics, and more. Metabolic engineering, the use of microbes to produce small and large molecules of commercial and scientific interest, has been a particular focus. To this end, synthetic biologists have developed methods to control transcription and translation, knock out native genes to route metabolic flux down desired channels, integrate multiple chemical and physical inputs to make decisions inside microbial cells, and make wholesale changes to the genomes of organisms in high throughput.

A new paradigm in synthetic biology technologies focuses on a new challenge: spatiotemporal organizational of biochemical processes. Organisms in all domains of life exert fine control over when and where biochemical reactions occur, be they responsible for metabolism, information transfer, or cell replication. This kind of organization remains conspicuously absent from most engineered systems.

Early efforts in synthetic biology focused on the creation of generalized systems to transcribe and translate heterologous genes and proteins in a variety of hosts, from bacteria to fungi to mammalian cells. These included the creation of modular libraries of genetic parts, such as plasmids, promoters, terminators, and ribosome binding sites (Beal et al., 2014; Blazeck et al., 2012b, 2012a; Brophy and Voigt, 2016; Dahl et al., 2013; Fernandez-Rodriguez and Voigt, 2016; Leavitt et al., 2016; Lee et al., 2013, 2015; Mutalik et al., 2013; Rajkumar et al., 2016; Salis et al., 2009). It is now possible to computationally design composable genetic circuits based on these parts with high fidelity (Nielsen et al., 2016; Roehner et al., 2016). Efforts were also made to understand and control translation elongation, and inform the choice of codons in heterologous genes (Goodman et al., 2013).

Along with the introduction of foreign genes into microbial factories, it was soon recognized that removing host genes was also of critical importance. Sophisticated computational approaches now exist to predict which native genes should be removed from a microbe in order to maximize the yield of a desired biosynthetic product (Burgard et al., 2003; Price et al., 2004; Alper et al., 2005; Kim et al., 2008). These models can also include predictions of host or foreign genes whose introduction might be beneficial (Ranganathan et al., 2010; Srivastava et al., 2012).

Cellular computation, enabling cells to collect information from their environment and make decisions accordingly, has also been an important focus of synthetic biology (Purcell and Lu, 2014), but will not be discussed further here. Instead we will focus on metabolic engineering applications.

Despite all of these technological advances, there have been relatively few examples of the commercially successful, industrial scale production of chemicals by microbes (Lechner et al., 2016). Notable successes have included artemisinic acid (Paddon et al., 2013), as well as farnesene, 1,3-propanediol, and 1,4-butanediol (Lechner et al., 2016), but, by and large, the biological production of chemicals at industrially viable titers has remained elusive. This is most often due to one (or more) of five ubiquitous roadblocks to biosynthesis: cellular toxicity due to accumulation of intermediates of the biosynthetic pathway; the loss of flux to undesired byproducts; difficulties sustaining sufficient substrate influx; leakage and loss of intermediates into the culture medium; and trapping of product in the host cell due to inadequate efflux [Fig. 1A].

**Figure 1.**
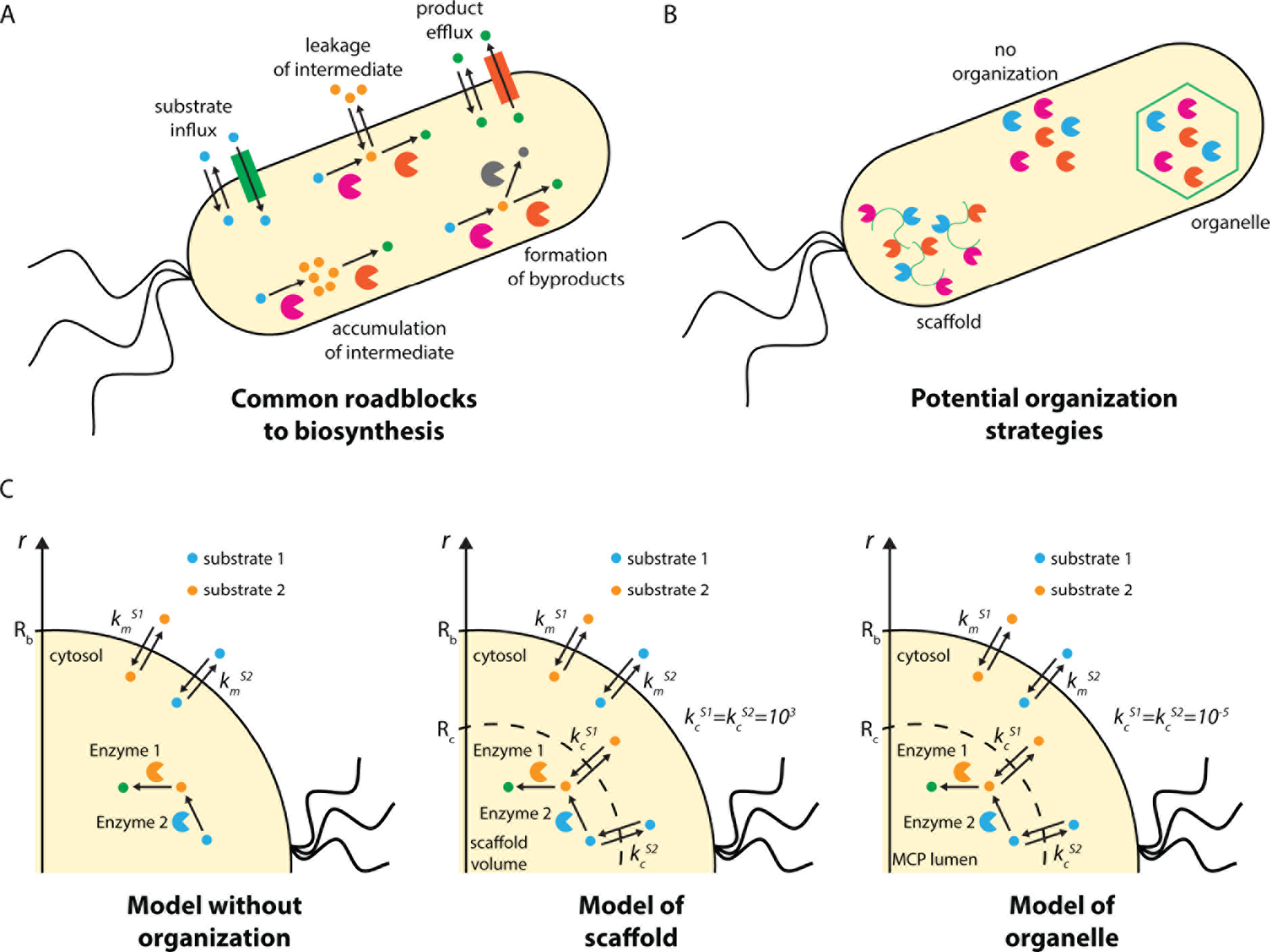
(A) Roadblocks commonly facing heterologous biosynthesis. (B) Potential organization strategies to alleviate these roadblocks. (C) Schematics of our models of (left) a pathway without organization, (middle) a pathway organized on a scaffold, and (right) a pathway organized in an organelle.

A new wave of synthetic biology technologies aims to address these key issues using a diverse array of strategies, while also preparing to deploy engineered microbes widely and safely. These cutting-edge approaches include cell-free approaches to protein and small molecule synthesis (Dudley et al., 2016; Garamella et al., 2016; Goering et al., 2016; Lu et al., 2015, 2014; Sullivan et al., 2016; Worst et al., 2015), dynamic control of metabolite concentrations (Xu et al., 2014) and of transcription and translation at the RNA level (Chappell et al., 2015; Takahashi and Lucks, 2013), robust approaches to biocontainment (Lopez and Anderson, 2015; Mandell et al., 2015), sensing of diverse small molecules (Mukherjee et al., 2015), establishing consortia of synergistic microbes (Marchand and Collins, 2016; Peng et al., 2016), and discovering enzymes facilitating previously unknown catalyses (Walker et al., 2013; Zhu et al., 2015). Broadly speaking, these strategies address a key missing capability in the synthetic biological toolkit: the ability to control precisely when and where chemical reactions take place (Boyle and Silver, 2012; Kerfeld, 2017; Kim and Tullman-Ercek, 2013).

Here, we will analyze one class of these strategies: the spatial organization of metabolism within cells [Fig. 1B] (Polka et al., 2016). There are a wide variety of natural methods for spatial organization (Agapakis et al., 2012). Eukarya discretize their biochemistry into highly chemically distinct subcellular compartments and into enzyme complexes such as polyketide synthases (Khosla et al., 2014) and other metabolons (Wu and Minteer, 2014). Bacteria, too, are now understood to organize their metabolism in a variety of ways, including using protein-based carboxysomes (Shively et al., 1973), microcompartments (Bobik et al., 1999), and encapsulins (McHugh et al., 2014). 1,2-propanediol utilization (Pdu) microcompartments, for instance, are protein-bound organelles of approximately 150 nm diameter. The enclosing protein shell consists of trimeric, pentameric, and hexameric protein tiles with central pores that permit the passage of small molecules in and out of the organelles, but which prohibit the passage of enzymes and other proteins. Building on foundational microbiological understanding of these systems (Bobik et al., 1999; Fan et al., 2010; Huseby and Roth, 2013; Kerfeld et al., 2005; Kofoid et al., 1999), we and others have demonstrated control of the formation (Kim et al., 2014), protein content (Jakobson et al., 2015; Lawrence et al., 2014; Parsons et al., 2010; Wagner et al., 2016), catalytic activity (Jakobson et al., 2016), and transport properties (Slininger Lee et al., 2017) of these organelles. Due to their relative simplicity and ease of manipulation, these various protein-based compartments make excellent model systems for exploring the role of spatial organization on metabolism.

Likewise, much work has been done to characterize carboxysomes—compartments in which CO_2_ is concentrated to enhance carboxylation in photosynthetic bacteria-- and adapt them for engineering purposes (Cai et al., 2015, 2016; Chen et al., 2013). The modularity of carboxysomes and other CO_2_ concentrating mechanism components facilitates reconstitution in other organisms (Bonacci et al., 2012; Gonzalez-Esquer et al., 2015). An active area of research is reconstitution in plants (Lin et al., 2014a, 2014b; Long et al., 2015), as part of a broad strategy to increase plant yields (Giessen and Silver, 2017; Hanson et al., 2016; Rae et al., 2017; Sharwood et al., 2016). Engineering microbial metabolism with CO_2_ as the primary carbon source would allow the production of sustainably produced biofuels and other high value products (Antonovsky et al., 2016; Ducat and Silver, 2012) and carboxysomes could enhance such strategies. More generally, ccarboxysomes have been suggested as modular method for partitioning nonnative pathways from native metabolism (Kerfeld, 2017). However, we have strong indication that performance of CO_2_ concentrating mechanisms will depend on how encapsulation interplays with transporters or other exogenous conditions setting the supply of CO_2_ (Mangan and Brenner, 2014; Mangan et al., 2016), and further systems analysis is required to realize the benefits of carboxysome-based encapsulation (Hanson et al., 2016; Long et al., 2016).

Bacterial microcompartment organelles are not the only organization solution available to the metabolic engineer; scaffolds based on protein, lipid, DNA, and RNA have all shown promise in improving heterologous pathway performance. Each of these strategies have been shown to be effective for the enhancement of heterologous biosynthesis in various contexts (Conrado et al., 2012; Delebecque et al., 2011; Dueber et al., 2009; Lawrence et al., 2014; Moon et al., 2010; Myhrvold et al., 2016). These studies organized diverse biosyntheses, including of mevalonate, resveratrol, 1,2-propanediol, and molecular hydrogen, suggesting that many different enzymatic pathways could be enhanced by scaffolding.

Having developed tools to control the localization of heterologous biosynthetic pathways to these organizing structures, a crucial question remains: what pathways are suitable for organization? And what benefits might be accrued by organizing pathways in one way versus another?

## Results

### Pathway encapsulation in bacterial microcompartments can provide benefits comparable to protein engineering

We will outline the potential metabolic engineering benefits that could be derived from pathway organization using two different strategies: encapsulation in the Pdu microcompartment of *Salmonella* and other enteric bacteria, and organization using a scaffold (which could be organized by means of protein, lipid, or nucleic acid) [Fig. 1C].

To address the kinetic consequences of encapsulation in these structures, we make use of a computational framework (Jakobson et al., 2017) developed to analyze the native function of the microcompartment organelles. A key prediction of this model is that microcompartments significantly enhance pathway flux. We predict that the natively encapsulated system enjoys a four-order of magnitude enhancement in flux upon encapsulation, as compared to free diffusion of the enzymes in the cytosol (Jakobson et al., 2017). Appropriately chosen heterologous pathways might also accrue such flux enhancements, as well as potentially reducing the loss of pathway intermediates to the extracellular space.

To instead model a scaffolded system, we simply assume that the enzymes in question are localized to a volume equivalent to that occupied by the Pdu microcompartments, but without a diffusion barrier. This is represented mathematically by setting the velocity of transport between the scaffold volume and the cytosol equal to that predicted by free diffusion. The model formulation we use here is agnostic to the underlying scaffolding platform (protein, lipid, or nucleic acid) or its microscopic organization, as we assume a well-mixed scaffold volume.

We predict the consequences of organization using these two strategies for two model biochemical processes: native Pdu microcompartment metabolism and the heterologous mevalonate biosynthetic pathway. The compounds and enzyme kinetic parameters for each pathway are in Figure 2AB. We adapt the modeling approach used for the native Pdu system to make flux predictions for the heterologous mevalonate pathway by adjusting the enzymatic kinetic parameters, cell membrane permeability to metabolites, and enzyme abundance and stoichiometry. While the first substrate of the mevalonate pathway (acetoacetyl-CoA) is produced intracellularly, rather than entering from the extracellular space (in the case of 1,2-PD), we approximate the generation of acetoacetyl-CoA upstream as a constant extracellular concentration in the context of our kinetic model. This approximation could correspond to production of acetoacetyl-CoA by a relatively faster and reversible upstream enzyme, or to more complex homeostatic control of the acetoacetyl-CoA concentration in the cytosol, both resulting in an effectively constant concentration of acetoacetyl-CoA far from the organelle or scaffold. Supporting this assumption, we find that the concentration gradient in the cytosol is small across a wide range of external substrate concentrations for all the organizational cases we tested (Fig. S1).

**Figure 2.**
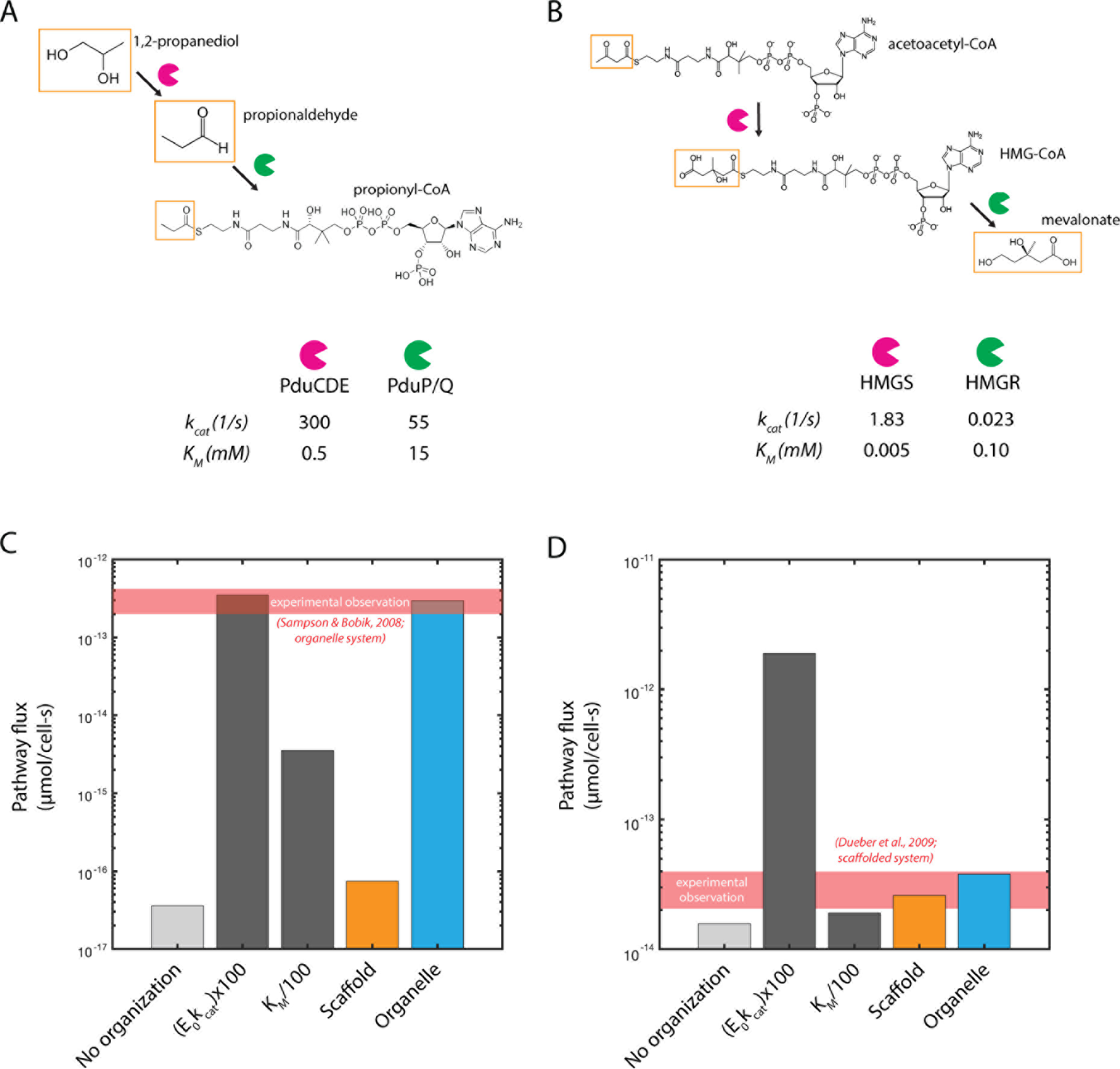
Relevant substrates, intermediates, products, and enzyme kinetic parameters for (A) native Pdu microcompartment metabolism and (B) mevalonate biosynthesis. Predicted pathway flux for (C) native Pdu microcompartment metabolism and (D) mevalonate biosynthesis for native kinetics without organization; 100-fold improvement of *k*_*cat*_ for both pathway enzymes; 100-fold improvement in *K*_*M*_ for both pathway enzymes; native kinetics with organization on a scaffold; and native kinetics with organization in a microcompartment organelle. The predictions here are based on an external substrate concentration of 50 mM 1,2-propanediol, as is typically used in experiments (Sampson and Bobik, 2008). We use the same external substrate concentration (50 mM) in the mevalonate case. Experimental observations in (C) and (D) are calculated from *S. enterica* growth rates from Sampson and Bobik, 2008 and from titer measurements for a scaffolded system from Dueber et al., 2009.

**Figure 3.**
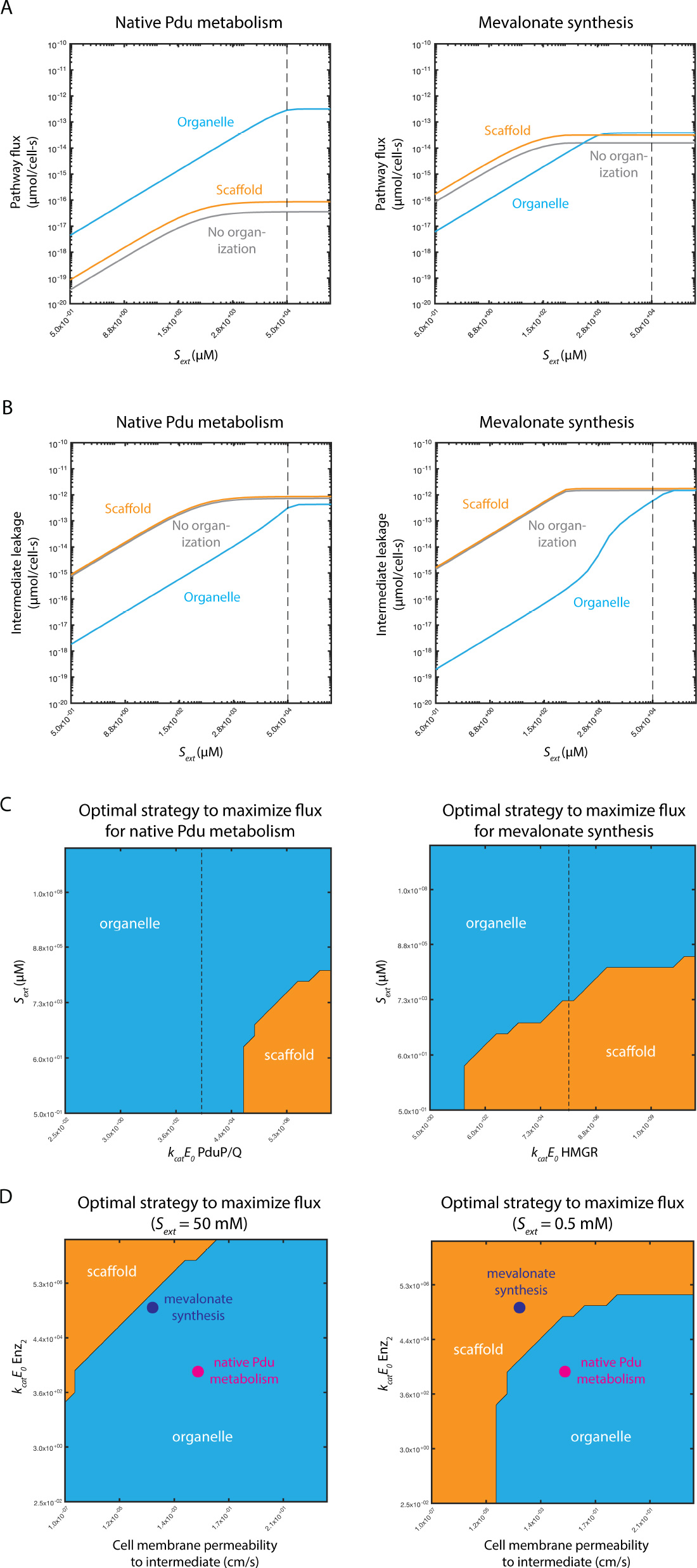
(A) Predicted flux for (left) native Pdu microcompartment metabolism and (right) mevalonate biosynthesis without organization (grey); with organization on a scaffold (orange); and with organization in an organelle (blue) as a function of external substrate concentration *S*_*ext*_. (B) Predicted intermediate leakage for (left) native Pdu microcompartment metabolism and (right) mevalonate biosynthesis without organization (grey); with organization on a scaffold (orange); and with organization in an organelle (blue) as a function of external substrate concentration *S*_*ext*_. (C) Predicted optimal organizational strategy for (left) native Pdu microcompartment metabolism and (right) mevalonate biosynthesis as a function of external substrate concentration *S*_*ext*_ and the abundance and kinetics of the second pathway enzyme *k*_*cat*_*E*_0_ (PduP/Q and HMGR, respectively). Baseline parameter values are shown with a black dashed line. (D) Predicted optimal organizational strategy for native Pdu microcompartment metabolism (magenta) and mevalonate biosynthesis (purple) as a function of the abundance and kinetics of the second pathway enzyme *k*_*cat*_*E*_0_ (PduP/Q and HMGR, respectively) and the cell membrane permeability to the intermediate at (left) external substrate concentration *S*_*ext*_ = 50 mM and (right) *S*_*ext*_ = 0.5 mM. Regions of parameter space are colored by the optimal organization strategy in that region: organelle (blue); scaffold (orange); or no organization (grey).

We first ask: is organization worth the time and trouble for the metabolic engineer to arrange, as compared to traditional metabolic and enzyme engineering strategies (such as improvements to the *k*_*cat*_ or *K*_*M*_ kinetic parameters)? Here, we use the native Pdu microcompartment pathway as an example. If, for instance, engineering efforts increased the *k*_*cat*_ of each of the two key enzymatic steps 100-fold, or decrease the *K*_*M*_ of each key step 100-fold, the improvement in flux would be as shown in Figure 2CD (as compared to the native system with no organization). Improvements of this magnitude for both enzymes represent a significant technical challenge and would be non-trivial to achieve for an arbitrary enzymatic system. We predict that encapsulation of native Pdu metabolism in an organelle is practically as effective as large improvements in *kcat*, and more effective than large improvements in *K*_*M*_, with respect to increasing the total flux through the pathway [Fig. 2C]. See the Methods for a detailed description of this calculation and the model in general. The predicted concentrations of each metabolite for each kinetic case are shown in Figure S2. The dramatic improvement in predicted flux upon encapsulation is due to a large increase in the intermediate concentration in the vicinity of the second pathway enzyme, exceeding the saturating concentration. Moreover, this benefit comes without a significant increase in the cytosolic concentration of this intermediate [Fig. S2]. Our simulations predict the native Pdu microcompartment metabolic pathway benefits substantially more from encapsulation than it would from scaffolding [Fig. 2C].

On the other hand, the pathway to produce mevalonate accrues similar flux enhancement from an organelle- or scaffold-based organization strategy [Fig. 2D; Fig. S3]. Our prediction agrees with the experimental observation that organizing the mevalonate pathway on a protein scaffold increased titers (Dueber et al., 2009). We predict marginal additional benefit from an encapsulation approach in this case, since the Michaelis-Menten constants *K*_*M*_ for the enzymes are small, whereas we predict the potential for flux enhancement by encapsulation in an organelle is high for pathways kinetically similar to native Pdu metabolism (that is, with larger *K*_*M*_) [Fig. 2CD; see also Fig. 4B]. These kinds of predictions can be made *a priori* for any enzymatic pathway for which the kinetic parameters are known (or can be approximated).

**Figure 4.**
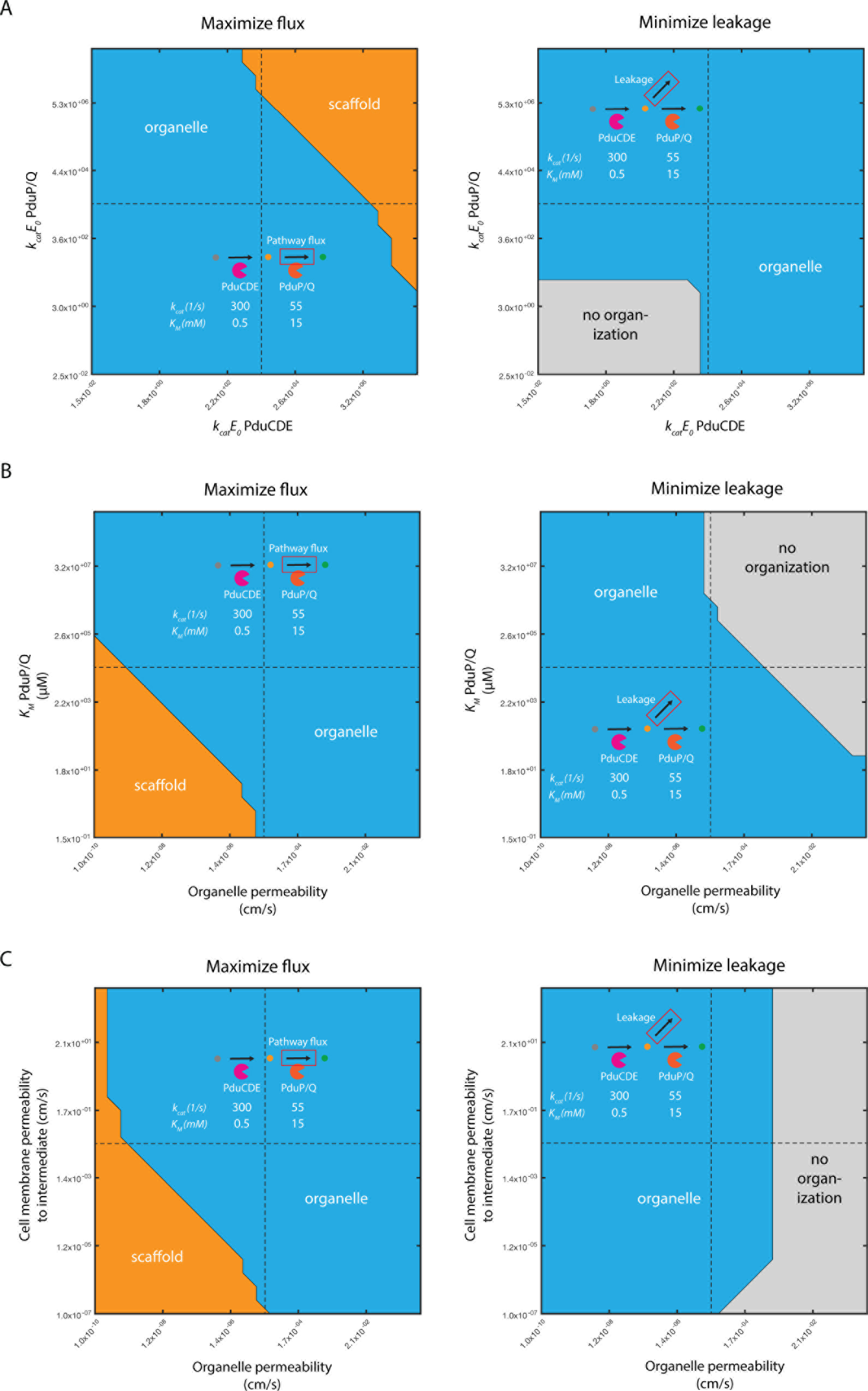
Recommended organization strategy resulting in (left) maximum pathway flux or (right) minimum intermediate leakage for native Pdu metabolism as a function of (A) *k*_*cat*_*E*_0_ of PduCDE and PduP/Q, (B) organelle permeability and *K*_*M*_ of PduP/Q, and (C) organelle permeability and cell membrane permeability. Regions of parameter space are colored by the optimal organization strategy in that region: organelle (blue); scaffold (orange); or no organization (grey). Baseline parameter values are shown with a black dashed line.

In both cases, the increased pathway flux in the case of increasing the *k*_*cat*_ of each enzymatic step comes at the cost of greatly increased loss of intermediate species to the extracellular space (or to other cellular process, in the case that there are competing reactions in the cytosol) [Figure S4]. This tradeoff may be important to consider in some cases, for instance if the intermediate species is toxic, and may render the protein engineering strategy less appealing than organization, despite similar predicted flux enhancement.

### Optimal organization strategies for biosynthetic pathways differ based on pathway properties and culture conditions

In addition to intrinsic properties of the pathway in question, the benefits of encapsulation versus scaffolding can depend on extrinsic factors, such as the bulk concentration of substrate. At lower external substrate concentrations, flux for pathways organized with scaffolds improves relative to an organelle for both pathways we considered [Fig. 3A]. This is because the rate of entry of substrate into the organelle becomes problematic at low bulk substrate concentrations, when the driving force for transport into the organelle is reduced. The rate of leakage of intermediate to the extracellular space is also affected; in each case, a scaffold leads to the greatest intermediate leakage, and this disadvantage worsens at low bulk substrate concentration for both pathways [Fig. 3B]. This underscores the importance of considering the pathway in question and the desired outcome (flux enhancement or leakage prevention) when selecting organization strategies. Modeling approaches could be extended in future to account for this duality by creating composite objective functions for the energetic cost of flux enhancement and leakage, or for the cost of the enzymes and organizing structures themselves (Noor et al., 2016), and optimizing across these different factors simultaneously.

We next consider the effects of one cell-intrinsic property (the abundance and kinetics of the second pathway enzyme) and one cell-extrinsic property (external substrate concentration) simultaneously [Fig. 3C]. These phase spaces show the optimal organizational strategy to maximize flux as a function of both variables, with the optimal strategy indicated by the color of the phase space at that parameter value combination. For reference, the dashed line in each panel of Figure 3C indicates the *k*_*cat*_*E*_*0*_ value used to construct the one-dimensional representations with respect to *S*_*ext*_ in Figure 3A. While the topology of these landscapes is similar for both systems (and indeed for any irreversible two-enzyme pair governed by Michaelis-Menten kinetics), there are important quantitative differences. Critically, given the *k*_*cat*_*E*_*0*_ value of the second enzyme in the mevalonate synthesis pathway, we predict that organelle-type organization is favored at high *S*_*ext*_ but scaffolding is favored at low *S*_*ext*_ values [Fig. 3C]. A batch-type reactor, therefore, might transition from organelles to scaffolds being optimal during a production run; laboratory-scale pilot experiments are most often conducted in this mode, potentially convoluting different optimality regimes. This observation holds for *k*_*cat*_*E*_*0*_ values several orders of magnitude smaller or larger than our estimate. The same is not true for the native Pdu MCP system, in which organelles are favored for all *S*_*ext*_ values given our assumptions regarding *k*_*cat*_*E*_*0*_ of PduP/Q [Fig. 3C]. This kind of information is key in designing optimally productive biosynthetic processes, and might call for a dynamic organizational transition as culture conditions change (Yang et al., 2017).

Finally, we demonstrate how the optimal organizational strategy changes as a function of two intrinsic pathway properties (rather than one intrinsic and one extrinsic property, as in Fig. 3C). While there are many parameters that can change between pathways, we focus on two key differences between the Pdu MCP system and mevalonate pathway. The values we estimate for the cell membrane permeability to the intermediate and *k*_*cat*_*E*_*0*_ for the second enzyme differ by approximately two orders of magnitude between the two systems. We therefore predicted the optimal organizational strategy as a function of these two parameters, and indicated the location of each enzyme system in this phase space [Fig. 3D]. We set all model parameters besides cell membrane permeability and *k*_*cat*_*E*_*0*_ to the baseline values for the Pdu MCP system. The phase space does not qualitatively change if we instead set all the other parameters to those representative of the mevalonate biosynthetic pathway [Fig. S5]. We constructed the phase space for two external substrate concentrations *S*_*ext*_, 50 mM and 0.5 mM. Crucially, the change in external concentration shifts the boundary between the regions in which scaffold and organelle strategies are optimal. At the lower *S*_*ext*_, mevalonate biosynthesis favors a scaffold over an organelle, while organelle organization is still favored for the Pdu MCP.

Phase spaces of the kind we explore here can be constructed for any pair of parameters, and provide a means to survey the organizational performance landscape comprehensively across a very wide range of possible parameter values. The parameters to explore could include those susceptible to manipulation via culture conditions, such as *S*_*ext*_; those that can in principle be engineered, such as *k*_*cat*_*E*_*0*_; and those that are intrinsic to the relevant biomolecules, such as the cell membrane permeability to the intermediate species. By constructing these phase spaces for a variety of parameters, the metabolic engineer can gain a quantitative understanding of which parameters are critical in determining the optimal organization strategy for a given pathway, and can weigh the ease of altering a given parameter against the potential rewards in terms of engineering goals like pathway flux.

### Enzyme stoichiometry and kinetics, as well as design goals, influence optimal organization strategy

We can address the question of organization choice more generally by examining the relative performance of three strategies (free cytosolic localization of enzymes; scaffolds; and microcompartments) across a wide range of enzyme kinetic parameters. Considering, for instance, the two-enzyme pathway of native Pdu metabolism, we can predict the optimal strategy as we vary the activity (*k*_*cat*_*E*_*0*_) of each enzyme [Fig. 4A]. This variation can represent either the Pdu enzymes or a different enzymatic pathway. In this example, microcompartment organization is favored with respect to maximizing pathway flux unless the respective *k*_*cat*_*E*_*0*_ parameters for both enzymes are sufficiently large to render the effect of concentrating intermediate species in the microcompartment insignificant, in which case a scaffold is recommended. Conversely, for small *k*_*cat*_*E*_*0*_ of both enzymes, a strategy without spatial organization is indicated to minimize intermediate loss [Fig. 4A]. These phase space predictions of organization performance can be made for arbitrary organizational strategies and metabolic pathways, given appropriate kinetic models.

### The chemical character of the substrate and intermediate also have an impact on organization choice

In addition to the kinetic properties of the pathway enzymes, we can consider the effect of different substrates and intermediates on the choice of appropriate organizational strategies. Once again we compute the recommended organization strategy for the native Pdu metabolic pathway, and vary the values of the relevant model parameters. In this case, several parameters could be affected by the chemical character of the species, including transport across the cell membrane and transport in and out of the microcompartment organelle. If, for instance, transport of the substrate and intermediate across the microcompartment shell is slow, scaffold expression may be favored if the *K*_*M*_ of the second enzyme is sufficiently low [Fig. 4B]. On the other hand, if escape of the intermediate across the cell membrane is slow, scaffolding may be favored regardless of enzyme kinetics [Fig. 4C], since the cell itself can perform the organelle’s intermediate-concentrating function in this case. It may be possible to optimize the permeability of the microcompartment shell to the kinetics of the desired pathway and broaden the range of conditions under which an organelle is the optimal pathway (Park et al., 2017; Slininger Lee et al., 2017). All of these factors must be considered when choosing an appropriate organization strategy. This is particularly important when comparing biosynthetic pathways with substrates and intermediates of different sizes, which might reasonably be expected to have disparate transport properties at the cell membrane and microcompartment shell.

### Enzyme mechanism can alter the potential benefits of organization strategies

In the above examples, we consider irreversible, Menten-Michaelis kinetics for each enzymatic step of each pathway. This assumption holds for the Pdu microcompartment case, but not for all systems. For example, in the carboxysome, a carbon-fixation organelle of cyanobacteria, a key enzymatic step catalyzing the interconversion of CO_2_/HCO_3_^−^ is reversible, limiting the benefit of organelles to concentrate intermediate species (Mangan and Brenner, 2014). Selective permeability of the carboxysome does not result in increased CO_2_ concentration (Mangan et al., 2016). The reversibility of the CO_2_/HCO_3_^−^ conversion imposes a fundamental limit on the concentration of CO_2_ that can be achieved in the organelle [Fig. S6A]; in the microcompartment, on the other hand, selective permeability combined with enzyme irreversibility allows the development of a very high local intermediate concentration if the intermediate is selectively trapped [Fig. S6B; purple line]. The comparison between the reversible and irreversible kinetic models highlights the need to account for detailed aspects of kinetic mechanism, such as cofactors, inhibition, and other dynamic effects. Recent studies have also indicated that the local chemical environment of nucleic acid-based scaffolds can have beneficial effects on enzyme kinetics (Zhang et al., 2016); detailed kinetic effects of this kind can be incorporated into future models as they are elucidated.

## Discussion

### A fundamental framework for organizational choice

Above, we outline the potential for the spatial organization of heterologous pathways to greatly enhance their performance. This approach compares well with traditional enzyme engineering approaches. Furthermore, we describe a general framework to guide the choice of appropriate organizational strategies for metabolic engineering. Several key parameters must be accounted for: (I) enzyme kinetics; (II) substrate and intermediate chemical properties; and (III) external culture conditions. Some of these properties, such as the external substrate concentration and the transport properties of the cell membrane, can influence the supply of substrate to the pathway, while others, such as the presence or absence of competing reactions and the transport properties of the organelle boundary, can influence the loss of metabolic flux to off-target species. We outline the use of a general modeling approach to analyze the performance of different organization strategies, and present example organizational recommendations. Moreover, modeling approaches can suggest key experiments (*e.g.* variations of external substrate concentration) that may reveal important discrepancies in the performance of different organization strategies. The MATLAB code used to generate the graphics in this manuscript is freely available on GitHub (URL TBD), and we encourage members of the metabolic engineering and synthetic biology communities to explore the organizational performance landscapes for their own systems of interest. We also welcome suggestions of other useful kinetic or organizational regimes to include in future versions of the model.

### Avenues to improve understanding and prediction of optimal spatial organization

The framework above can provide important insights into the choice of optimal organizational strategies for heterologous pathways, but several important aspects of pathway organization remain unexplored. These include product export, cell size and morphology, competitive reactions, and the detailed organization of the organelles or scaffolds within the host cell. The organization of organelles within cells has been investigated in the context of a constant cytosolic metabolite concentration (Hinzpeter et al., 2017), and future efforts could combine this and our approaches.

Our model as currently implemented can incorporate extensions to explore some of these areas, but some questions, notably the effect of competitive cellular reactions and reactions upstream of the organized process, will require the integration of our model with larger-scale metabolic models of host processes. Exploring the effect of the detailed subcellular localization of the organizing structures themselves will likewise require modifications to our current mathematical framework.

### Practical engineering considerations

Within the framework described here, we evaluate only the pathway flux and intermediate leakage predicted for each potential organization strategy, neglecting the difficulty associated with engineering a particular strategy. It may transpire that, for certain pathways, scaffolding proves challenging to implement due to incompatibilities between the requisite protein tags and the enzymes in question, or that the pores of microcompartment shells are fundamentally of low permeability to the substrates of other pathways. Practical experience with these issues will continue to inform the choice of appropriate strategies, and may eventually allow the integration of such practical as well as theoretical considerations into objective functions. We expect that some key experiments, such as assays to quantitatively determine the permeability of protein shells to small molecules, will greatly enhance the predictability of engineering pathways in microcompartments.

### Closing thoughts

Organizing biochemistry in both time and space holds tremendous potential to help deliver on the promise of synthetic biology: the ability to produce medically and industrially important molecules at high yield and high titer with minimal environmental disruption. Spatial organization of the kind we advocate is but one of many important approaches; techniques to use multiple (or no) cells, to detect and transport metabolites, and to exert dynamic control on short time scales are of critical importance, as well. We posit that detailed mechanistic models of each of these approaches will be key in building a fundamental theoretical understanding of the optimal strategies to improve the performance of arbitrary biosyntheses.

## Materials and Methods

### Contact for resource and reagent sharing

For questions and further information regarding software and models used herein, please contact NMM.

### Method details

#### Model

The reaction-diffusion model framework used herein is substantially the same as that described in Jakobson, *et al.*, PLoS Computational Biology, 2017, in which we explored the native function of the Pdu MCP system in *S. enterica*. The analytical and numerical approach is described in detail in that manuscript, but the important assumptions can be summarized as follows:

1. We assume a spherically symmetrical organelle or scaffold at the center of a spherically symmetrical cell.
2. We consider the system at steady-state.
3. We assume that the external concentrations of the substrate (*S*_*ext*_) and intermediate are constant.
4. We assume that the activity of the enzymes can be described by irreversible Michaelis-Menten kinetics.

The governing equations are as follows in the cytosol and in the organelle or scaffold-like structure:

Cytosol:

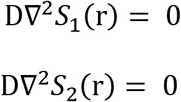

Organelle/scaffold:

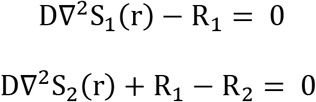

where

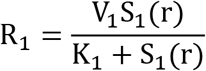

and

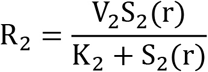

In the case of organelle- or scaffold-based organization, a closed-form analytical solution to the governing equations can be found, provided we assume that the metabolite concentration inside the organelle or scaffold region is constant. This solution is used to generate the various figures comparing organization strategies; see Jakobson et al., 2017 for the derivation and complete analytical solutions. A scaffold-like behavior is created by setting the permeability at the organelle boundary to approximate free diffusion (that is, k_c_^S1^ = k_c_^S2^ ~ 10^3^). In the case with no organization, we instead use a numerical solution, as the analytical solution does not hold in this regime. The numerical solution at steady state is generated by a finite difference routine implemented in MATLAB; again see Jakobson et al., 2017 for more details on the governing equations and boundary conditions used in the numerical routine.

#### Model parameters

The following table summarizes the important model parameters for the Pdu MCP system:

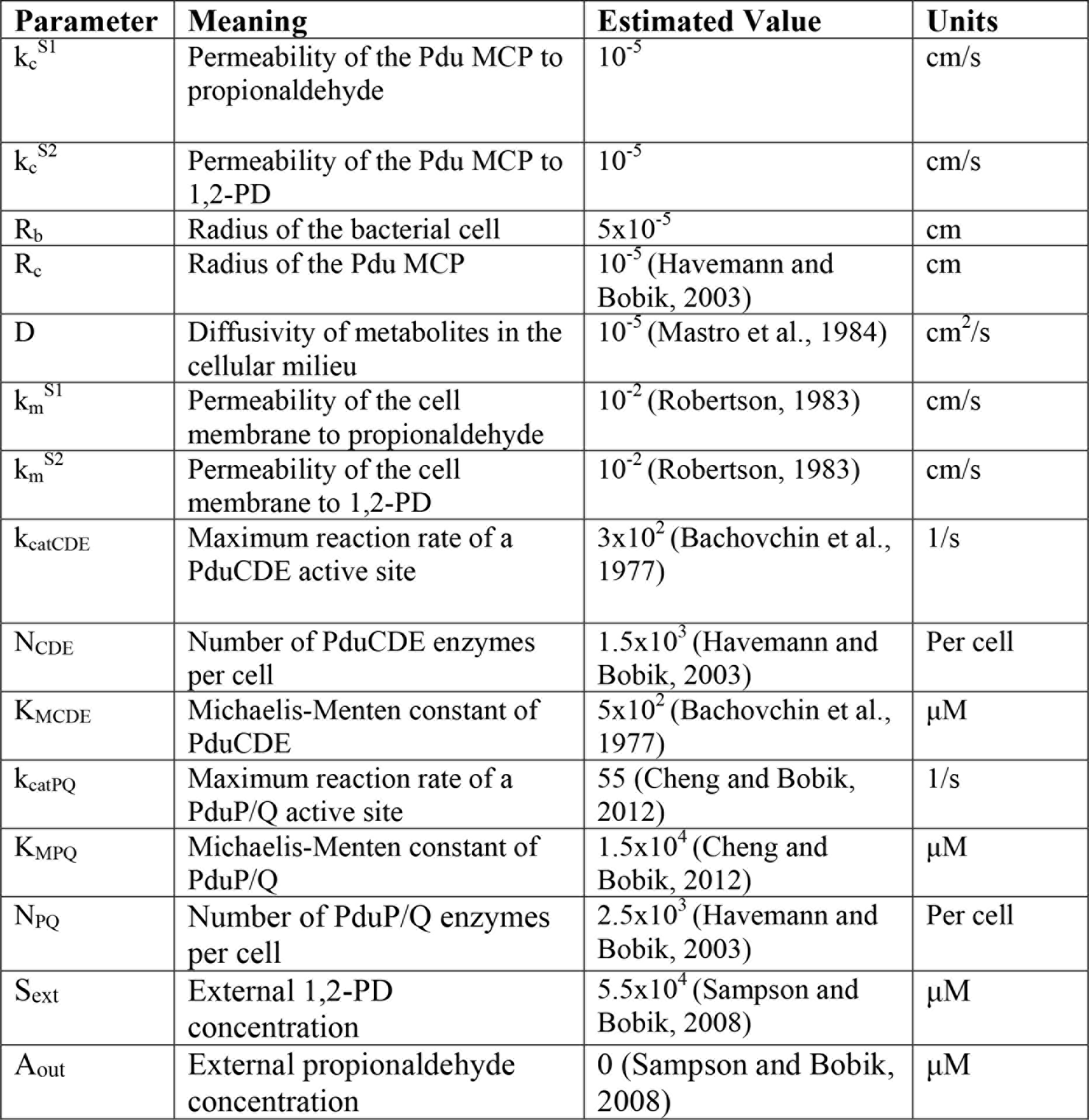

| Parameter | Meaning | Estimated Value | Units |
| --- | --- | --- | --- |
| $k_c^{S1}$ | Permeability of the Pdu MCP to propionaldehyde | $10^{-5}$ | cm/s |
| $k_c^{S2}$ | Permeability of the Pdu MCP to 1,2-PD | $10^{-5}$ | cm/s |
| $R_b$ | Radius of the bacterial cell | $5 \times 10^{-5}$ | cm |
| $R_c$ | Radius of the Pdu MCP | $10^{-5}$ (Havemann and Bobik, 2003) | cm |
| $D$ | Diffusivity of metabolites in the cellular milieu | $10^{-5}$ (Mastro et al., 1984) | $\text{cm}^2/\text{s}$ |
| $k_m^{S1}$ | Permeability of the cell membrane to propionaldehyde | $10^{-2}$ (Robertson, 1983) | cm/s |
| $k_m^{S2}$ | Permeability of the cell membrane to 1,2-PD | $10^{-2}$ (Robertson, 1983) | cm/s |
| $k_{\text{catCDE}}$ | Maximum reaction rate of a PduCDE active site | $3 \times 10^2$ (Bachovchin et al., 1977) | 1/s |
| $N_{\text{CDE}}$ | Number of PduCDE enzymes per cell | $1.5 \times 10^3$ (Havemann and Bobik, 2003) | Per cell |
| $K_{\text{MCDE}}$ | Michaelis-Menten constant of PduCDE | $5 \times 10^2$ (Bachovchin et al., 1977) | $\mu\text{M}$ |
| $k_{\text{catPQ}}$ | Maximum reaction rate of a PduP/Q active site | 55 (Cheng and Bobik, 2012) | 1/s |
| $K_{\text{MPQ}}$ | Michaelis-Menten constant of PduP/Q | $1.5 \times 10^4$ (Cheng and Bobik, 2012) | $\mu\text{M}$ |
| $N_{\text{PQ}}$ | Number of PduP/Q enzymes per cell | $2.5 \times 10^3$ (Havemann and Bobik, 2003) | Per cell |
| $S_{\text{ext}}$ | External 1,2-PD concentration | $5.5 \times 10^4$ (Sampson and Bobik, 2008) | $\mu\text{M}$ |
| $A_{\text{out}}$ | External propionaldehyde concentration | 0 (Sampson and Bobik, 2008) | $\mu\text{M}$ |

In the case of mevalonate synthesis, the parameters are altered as follows:

| Parameter | Meaning | Estimated Value | Units |
| --- | --- | --- | --- |
| $k_m^{S1}$ | Permeability of the cell membrane to acetoacetyl-CoA | $10^{-4}$ | cm/s |
| $k_m^{S2}$ | Permeability of the cell membrane to HMG-CoA | $10^{-4}$ | cm/s |
| $k_{catCDE}$ | Maximum reaction rate of a HMGS active site | 1.83 (Middleton, 1972) | 1/s |
| $N_{CDE}$ | Number of HMGS enzymes per cell | $5 \times 10^5$ (Dueber et al., 2009) | Per cell |
| $K_{MCDE}$ | Michaelis-Menten constant of HMGS | 5 (Cabañó et al., 1997) | $\mu M$ |
| $k_{catPQ}$ | Maximum reaction rate of a HMGR active site | 0.023 (Pak et al., 2008) | 1/s |
| $K_{MPQ}$ | Michaelis-Menten constant of HMGR | $1 \times 10^2$ (Pak et al., 2008) | $\mu M$ |
| $N_{PQ}$ | Number of HMGR enzymes per cell | $5 \times 10^5$ (Dueber et al., 2009) | Per cell |

#### Converting literature observations to flux predictions

For the systems we examine here, previous literature has described either cellular growth on 1,2-PD as the sole carbon source (Sampson and Bobik, 2008) or the volumetric titer of mevalonate in a scaffolded system (Dueber et al., 2009). To compare with our model, which generates predictions on a molecules-per-cell basis, we must convert these experimental observations to a comparable measurement.

For the native Pdu microcompartments, Sampson and Bobik observe a doubling time of approximately 5-10 hours during exponential phase growth on 1,2-PD as the sole carbon source (see Figure 3 of (Sampson and Bobik, 2008)). Assuming that a bacterial cell has a mass of approximately 0.3 pg and that half of the flux through the microcompartment pathway (by mass) can be used for cell growth, the Sampson growth observation predicts a steady-state flux of approximately 3×10^−13^ μmol/cell-s.

In the case of mevalonate biosynthesis by a scaffolded system, Dueber and colleagues report a titer of approximately 10 mM mevalonate after 2 days of culture, after which time the concentration changes little (see Figure 5 of (Dueber et al., 2009)). Assuming constant production over this time and a cell density of 2 OD~2×10^9^ cells/mL, this titer corresponds to a steady-state flux of approximately 3×10^−14^ μmol/cell-s.

### Quantification and statistical analysis

MATLAB R2016b (MathWorks) was used for all computation and to generate graphical representations of the results.

### Data and software availability

The model used herein is freely available on GitHub (URL TBD) under a GNU General Public License.

## Acknowledgments

This work was supported by the National Institutes of Health (grant number 1F32GM125162-01 to CMJ) and the National Science Foundation (award MCB1150567 to DTE). We thank the Tullman-Ercek and Jarosz laboratories for stimulating discussions.

The authors declare that they have no conflict of interest.

## Author contributions

CMJ, DTE, and NMM conceived of the project, analyzed results, and reviewed and edited the manuscript.

CMJ and NMM implemented the mathematical model and wrote the original draft of the manuscript.

DTE and NMM supervised the project.

